# Fission yeast Dis1 is an unconventional TOG/XMAP215 that induces microtubule catastrophe to drive chromosome pulling

**DOI:** 10.1101/2022.08.11.503693

**Authors:** Yuichi Murase, Masahiko Yamagishi, Naoyuki Okada, Mika Toya, Junichiro Yajima, Takahiro Hamada, Masamitsu Sato

## Abstract

The shortening of microtubules attached to kinetochores is the driving force of chromosome movement during cell division. Specific kinesins are believed to shorten microtubules but are dispensable for viability in yeast, implying the existence of additional factors responsible for microtubule shortening. Here, we demonstrate that Dis1, a TOG/XMAP215 ortholog in fission yeast, promotes microtubule shortening to carry attached chromosomes. Although TOG/XMAP215 orthologs are generally accepted as microtubule polymerases, Dis1 promoted microtubule catastrophe *in vitro* and *in vivo*. Notably, microtubule catastrophe was promoted when the tip was attached to kinetochores, as they steadily anchored Dis1 at the kinetochore-microtubule interface. Engineered Dis1 oligomers artificially tethered at a chromosome arm region induced the shortening of microtubules in contact, frequently pulling the chromosome arm towards the vicinity of spindle poles in meiocytes. Thus, unlike Alp14 and other TOG/XMAP215 orthologs, Dis1 plays an unconventional role in promoting microtubule catastrophe, thereby driving chromosome movement.

## Introduction

The dynamic behaviour of microtubules is essential for many aspects of cellular events, including chromosome segregation in dividing cells. Spindle microtubules capture the kinetochore region of chromosomes using the plus end and pull the chromosomes towards the spindle poles (centrosomes or yeast spindle pole bodies [SPBs]) by shortening the microtubules (also called depolymerisation) ^1^. For chromosome pulling in higher eukaryotes, depolymerisation occurs at the minus end of spindle microtubules around the poles. In contrast, depolymerisation is observed exclusively at the plus end, which is attached to kinetochores in yeast cells^2^.

By nature, microtubules demonstrate a dynamic property called dynamic instability *in vitro*, facilitating their spontaneous polymerisation and depolymerisation^3,4^. In cells, this property is further modulated by microtubule-associated proteins (MAPs), allowing microtubules to be transformed according to the cellular requirements.

Members of the kinesin-13 subfamily are microtubule catastrophe factors in higher eukaryotes, working at both ends of microtubules ^5^. However, kinesin-13 members are non-existent in the yeast genome, and kinesin-8 drives the depolymerisation of microtubules. Kip3, the budding yeast kinesin-8, depolymerises microtubules in a microtubule length-dependent manner ^6,7^. The kinesin-8 heterodimer Klp5/Klp6 in the fission yeast *Schizosaccharomyces pombe* is involved in microtubule destabilisation rather than depolymerisation ^8–11^. However, chromosome pulling could still be observed in Klp5/Klp6 knockout cells, indicating that other factors, possibly non-motor proteins, might be involved in microtubule shortening in fission yeast ^8,9,12,13^.

Members of the Dis1/TOG family are regulators of microtubule dynamics conserved across species. TOG has been accepted as a processive polymerase for microtubules, which provides tubulin dimers to the plus end ^14–19^. The *S. pombe* genome has two TOG orthologs, Alp14 and Dis1. Both Dis1 and Alp14 in fission yeast have been shown to locate at the plus end to polymerise microtubules ^20–23^. Alp14 was established as the major microtubule polymerase based on *in vitro* biochemical assays and the phenotype of Alp14 knockout cells. These cells exhibit short and fragile microtubules in the mitotic spindle, as well as in the cytoplasmic and radial arrays during interphase and meiosis ^21–24^.

Dis1, another TOG ortholog, reportedly promotes microtubule elongation in biochemical assays ^20^. However, the phenotype of Dis1 knockouts is puzzling: these cells display cytoplasmic microtubules of normal length similar to wild-type cells, and their dynamics are not largely altered ^25,26^. In mitosis, the spindle in Dis1 knockouts is fragile during prometaphase and relatively extended during anaphase ^11,27^. Furthermore, we have previously reported that chromosome pulling by microtubules was defective in *dis1Δ* meiocytes, indicating that Dis1 may depolymerise microtubule plus ends connected to kinetochores ^24^. Dis1 is also involved in shortening guanylyl-(αβ)-methylene-diphosphonate (GMPCPP)-stabilised microtubules *in vitro* ^20^. These findings suggest that Dis1 may play unconventional roles in shortening microtubules for chromosome pulling rather than canonical functions as a microtubule polymerase. Moreover, TOGs in other species may induce microtubule shortening according to circumstances ^28–34^.

Based on the increasing evidence, we propose that TOGs may have dual functions in extending or shortening microtubules and that Dis1 could be a previously unidentified candidate involved in microtubule shortening and drives chromosome pulling in concert with kinesin-8. Therefore, we investigated the possibility that Dis1 in fission yeast may shorten microtubules *in vitro* and *in vivo*.

## Results

### Dis1 induces microtubule catastrophe *in vitro*

In general, Dis1 (aka XMAP215 or TOG) family members are engaged in microtubule polymerisation, as evidenced by several *in vitro* and *in vivo* studies ^35^. Based on the growing evidence of the unique function of Dis1 in microtubule shortening, we examined the effect of Dis1 on microtubules *in vitro*.

Recombinant GST-Dis1 was incubated with fluorescein-labelled purified porcine tubulin, and microtubule formation was observed under the confocal microscope (Supplementary Fig. 1a). Microtubules in the presence of GST-Dis1 tended to shorten in a dose-dependent manner (Fig. 1a–c). GST-Dis1 also tended to decrease the number of microtubule bundles. However, this change was not statistically significant (Supplementary Fig. 1b). The decrease of bundles was not accompanied by a reduction in microtubule nucleation, implying that Dis1 does not inhibit nucleation (Supplementary Fig. 1c). The effect of Dis1 on microtubule shortening was further evaluated by the two-step incubation. First, microtubules were grown, and subsequently, GST-Dis1 was added prior to monitoring the microtubule dynamics (Fig. 1d; Supplementary Table 1). Microtubule growth was accelerated by GST-Dis1, whereas the shrinkage rate remained constant (Fig. 1e–g; Supplementary Table 1). Nonetheless, GST-Dis1 addition significantly increased the frequency of microtubule catastrophe (defined as an event in which growth turns into shrinkage) (Fig. 1e, h; Supplementary Table 1). The frequency of rescue (defined as an event in which shrinkage turns into growth) was also increased by GST-Dis1 (Fig. 1i; Supplementary Table 1). As a single catastrophe event quickly shortens microtubules in contrast to slow polymerisation, the net length of each microtubule was shortened in the presence of GST-Dis1 (Fig. 1b, c). Thus, Dis1 exerts its function directly on microtubules to induce catastrophe.

**Fig. 1:**
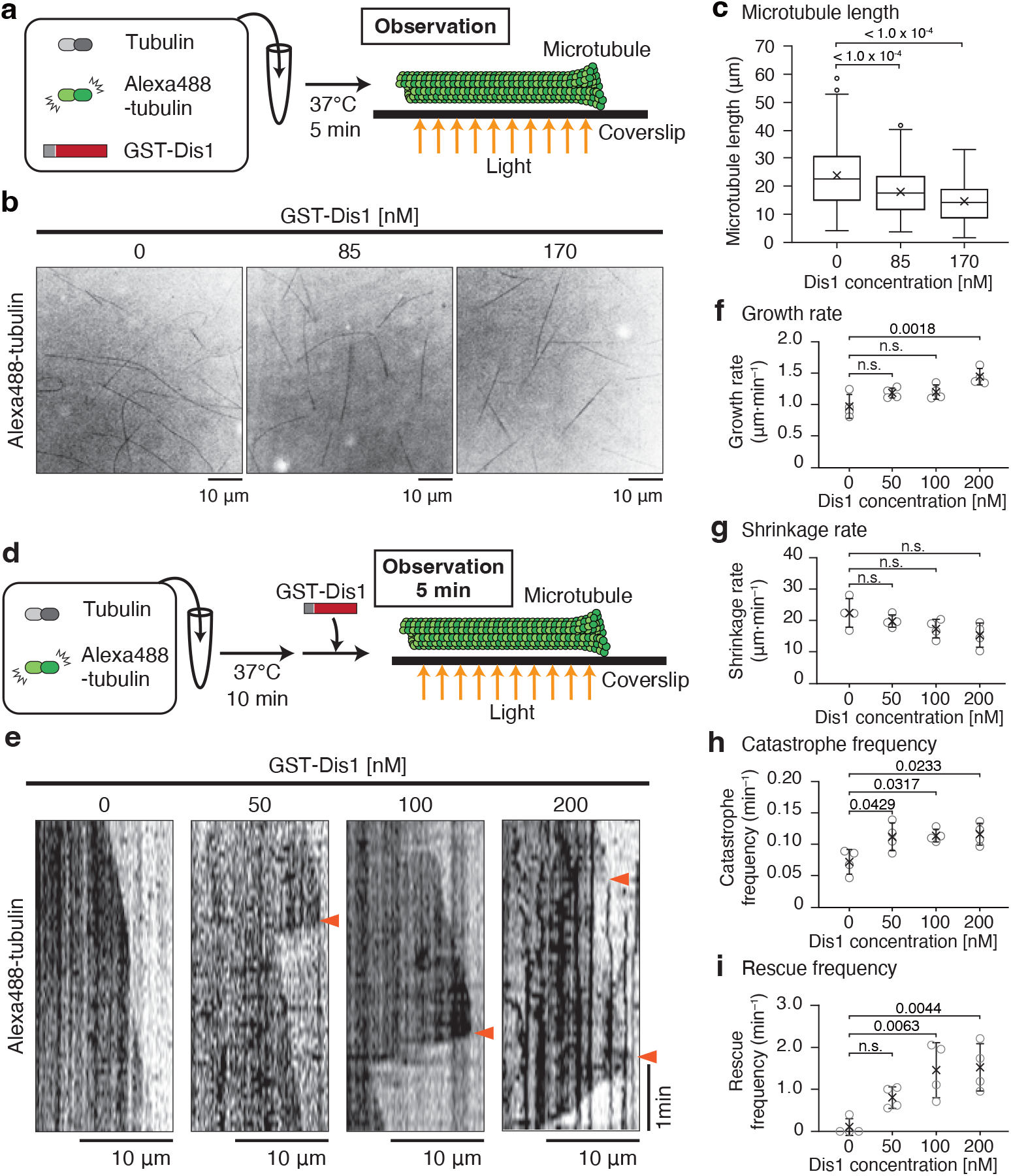
Dis1 induces microtubule catastrophes and shortens microtubules *in vitro*. **a** Experimental outline- purified tubulin (20 µM) with Alexa488-labelled tubulins (2µM) and 0, 85, and 170 nM of recombinant GST-Dis1 were mixed and incubated at 37°C for 5 min, followed by observation using a confocal microscope. **b** Observed microtubules with and without GST-Dis1. **c** Lengths of microtubules shown in (**b**) were plotted. Box-and-whisker plots indicate the minimum and maximum values and the 25^th^ and 75^th^ percentiles. Bullets indicate outliers; crosses represent means; Centre lines represent medians; *n* = 156 (0 nM), 197 (85 nM), 152 (170 nM) microtubules. **d** Experimental outline- purified tubulin (35 µM) with Alexa488-labelled tubulin (3 µM) was incubated at 37°C for 10 min. GST-Dis1 was then added and observed at 5-s intervals for 5 min using the fluorescent microscope. **e** Representative kymographs for Alexa488-labelled microtubules with indicated concentrations of GST-Dis1. Arrowheads represent the timing of microtubule catastrophes. **f**–**i** Growth (**f**), shrinkage rates (**g**), catastrophe (**h**), and rescue (**i**) frequencies were calculated. Crosses, the mean; bullets, technical replicates: *n* = 4 (0 nM), 4 (50 nM), 4 (100 nM), 4 (200 nM) experiments. At least 15 microtubules were observed for each experiment. Error bars; SD. The statistical significance of the difference was determined using one-way ANOVA followed by the Tukey–Kramer method. *P* values are shown; n.s., not significant.

### Dis1 anchored at kinetochores shortens microtubules in meiocytes

As previously demonstrated ^24^, the kinetochores of fission yeast are scattered in the nucleus during the meiotic prophase. They are then collected towards spindle poles at the onset of meiosis I by a radial array of microtubules emanating from the poles. In wild-type (WT) meiocytes, Dis1 was located at the plus-end of microtubules as well as kinetochores upon retrieval (Fig. 2a; Supplementary Fig. 2a). In *dis1Δ* (*dis1*-knockout) meiocytes, kinetochores were attached by the radial array of microtubules but were frequently uncollected (Fig. 2a). Collectively, these *in vitro* and *in vivo* results indicated that Dis1 might induce catastrophe, rather than depolymerisation, thereby shortening microtubules to retrieve kinetochores in WT meiocytes. Subsequently, catastrophe frequencies of microtubules in WT meiocytes were calculated with regard to Dis1 location. Microtubules decorated with (ii, Fig. 2b, c; Supplementary Table 2) and without (iii, Fig. 2b, c) Dis1-3GFP at the plus-end tended to undergo catastrophe to a similar degree, although the microtubules with Dis1-3GFP had a slightly higher catastrophe frequency. Notably, catastrophe was significantly promoted when the tip carried kinetochores concomitantly with Dis1 (i), implying that Dis1 and kinetochores synergistically promote catastrophe of the tip (*P* < 0.05, Fig. 2c; Supplementary Table 2). In contrast, the catastrophe frequency of microtubule tips in *dis1Δ* cells remained lower than in that in WT cells, irrespective of possession of kinetochores at the tip (iv, v; Fig. 2b, c; Supplementary Table 2), demonstrating that kinetochores do not inherently induce catastrophe in them associating microtubule tips.

**Fig. 2:**
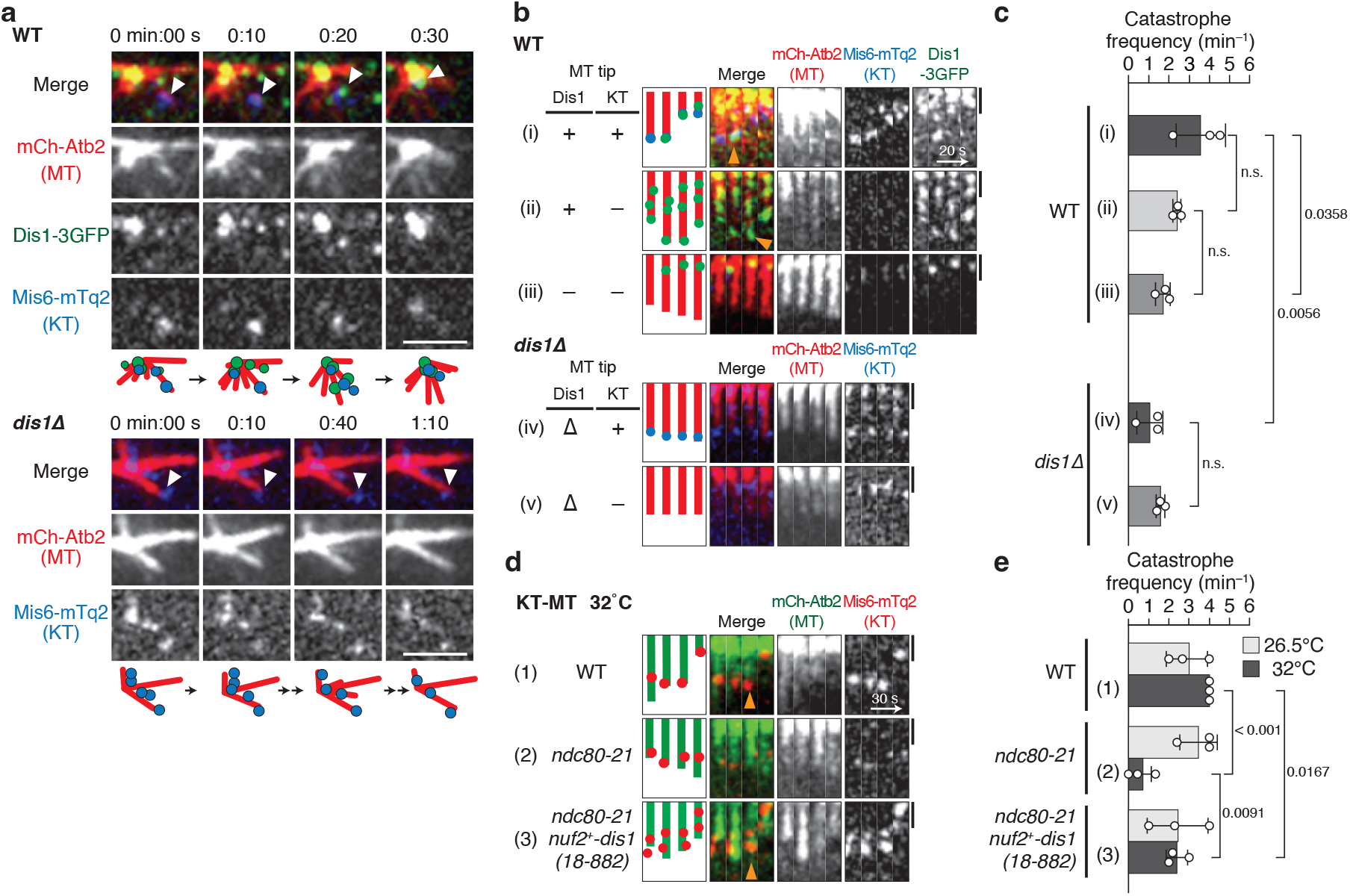
Dis1 induces microtubule catastrophe at the onset of meiosis I. **a** Time-lapse images of zygotic nuclei at the onset of meiosis I in wild-type (WT) and *dis1*Δ mutant cells. Dis1-3GFP (green) co-localises with kinetochores (KT; labelled with Mis6-mTurquoise2, blue) at microtubule tips (MT; mCherry-Atb2, red). The kinetochore-bound microtubule tip (arrowhead) started shortening at 0:20. Schematics are shown at the bottom. In *dis1*Δ, microtubules attached a kinetochore (arrowhead) but were not shortened. Scale bar, 2 µm. **b** Representative kymographs of microtubules classified by the state of the tips, shown with schematics. In WT, (i) tips with both kinetochores and Dis1; (ii) tips without kinetochores but with Dis1; (iii) tips without kinetochores or Dis1. In *dis1*Δ, with (iv) and without (v) kinetochores. The microtubules in (iv) are also shown in (**a**). Orange arrowheads represent microtubule catastrophes. Scale bar, 1 µm. **c** The catastrophe frequency for each category in (**b**); *n* = 3 (i), 3 (ii), 3 (iii), 3 (iv), 3 (v) experiments. **d** Representative kymographs for kinetochore-microtubules in (1) WT, (2) *ndc80-21* and (3) *ndc80-21 nuf2*^+^*-dis1(18-882)* cells at the onset of meiosis I at the restrictive temperature. Orange arrowheads represent microtubule catastrophe. Scale bar, 1 µm. **e** The catastrophe frequency for each state is shown in (**d**); *n* = 3 experiments for each sample. Bullets indicate technical replicates, and error bars indicate SD. The statistical significance of the difference was determined using one-way ANOVA followed by the Tukey–Kramer method. *P* values are shown; n.s., not significant.

To assess whether Dis1 induces frequent catastrophe, particularly when associated with kinetochores, we next employed the *ndc80-21* temperature-sensitive mutant. Ndc80/Hec1 is a component of the outer kinetochore complex ^36–38^, and Dis1 reportedly uses it as a platform to localise to kinetochores ^27^. Although Dis1 could localise to microtubules in *ndc80-21* mutant cells, the catastrophe frequency was decreased, as in *dis1*Δ cells (Fig. 2d, e; Supplementary Fig. 2b; Supplementary Table 3). The reduction was recovered by an enforced tethering of Dis1 to Nuf2 ^27^ (Fig. 2d, e; Supplementary Fig. 2b; Supplementary Table 3), another component of the Ndc80 complex ^39,40^. Collectively, we conclude that Dis1 actively induces microtubule catastrophe at the tip, particularly when located at kinetochores.

Thus, Dis1 in cells exerts its influence on the catastrophe of microtubule tips in concert with kinetochores. However, the mechanism by which kinetochores trigger the action of Dis1 remains unexplored. To this end, we investigated the behaviour of Dis1 at microtubule tips and observed that Dis1 tended to locate at microtubule tips for extended periods only when accompanied by a kinetochore than in its absence. In 77% of the microtubule shrinkage events we observed, Dis1 was detached from the tip when it did not accompany kinetochores (i, Fig. 3a; KT: –, Fig. 3b). The frequency of Dis1 that remained at kinetochores during shrinkage was significantly elevated when Dis1 accompanied a kinetochore at the tip (iii, Fig. 3a; 23% [KT: –] and 79% [KT: +] of the events, Fig. 3b). Dis1 tended to accumulate at the tip when attached to the kinetochores during shrinkage, whereas Dis1 unstably fluctuated in the absence of kinetochores (Fig. 3c). Dis1 does not stably locate at the plus tip of microtubules *in vivo* and *in vitro* (see Fig. 2b) ^20,26^. Therefore, we propose that kinetochores likely employ the Ndc80 complex ^27^ and serve as a platform to stably retain Dis1 at the shrinking microtubule tip.

**Fig. 3:**
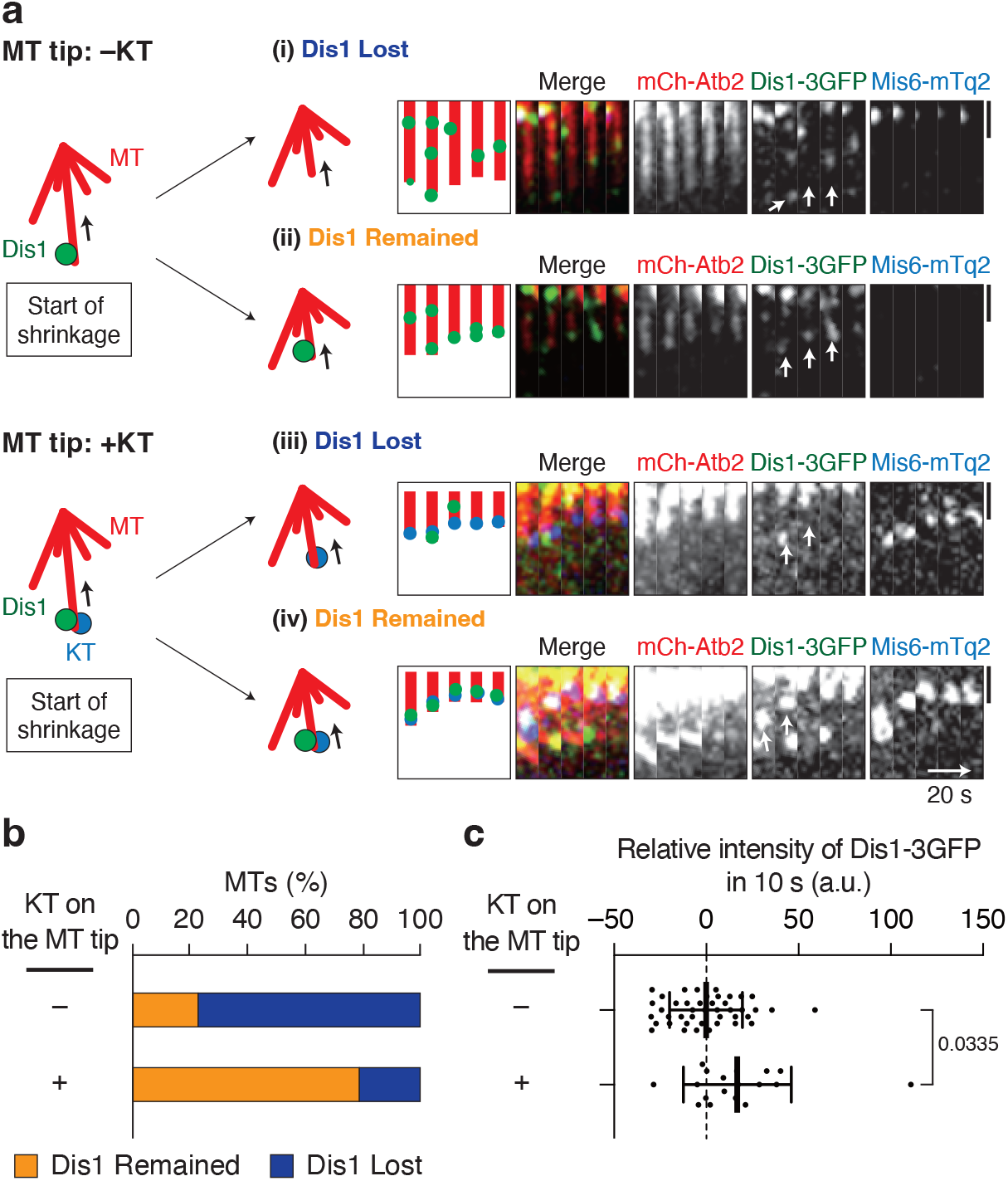
Kinetochore extends the duration of Dis1 location at the microtubule tip. **a** The fluorescence intensity of Dis1-3GFP localised at the tips of shrinking microtubules was measured in WT cells. Observed microtubules were classified into two groups: tips with Dis1-3GFP only (–KT, top) and tips with both Dis1-3GFP and kinetochores (+KT, bottom). Each group of microtubule tips was then observed over time to monitor the instability of Dis1-3GFP localisation at the tips: whether Dis1-3GFP was lost or decreased (‘Dis1 Lost’, i and iii), and alternatively, remained or increased (‘Dis1 remained’, ii and iv) during shrinkage. Representative kymographs and schematics are shown on the right. White arrows indicate tips of shrinking microtubules where the Dis1-3GFP intensity was measured. The microtubules in which the fluorescence intensities of Dis1-3GFP were lower than the initial value were classified as ‘Dis1 Lost’, and the rest was classified as microtubules with ‘Dis1 Remained’. **b** The rate of each event shown in (**a**) was quantified. *n* = 22 (–KT), 14 (+KT) microtubules. Dis1 was maintained at the end of microtubules when colocalised with kinetochore according to χ^2^ two-sample test (χ^2^ = 11, *P* < 0.005). **c** Fluctuation of Dis1-3GFP fluorescence intensity at the tip during shrinkage. The Dis1-3GFP intensity was measured at two time points of a 10-s interval, and the relative intensity of the second time point to the first was plotted. A plot above 0 means an increase of Dis1-3GFP in 10 s. Bold lines, means; error bars, SD; *n* = 43 (–KT), 18 (+KT) observations. The statistical significance of the difference was determined using Student’s two-tailed t-test; the *P* value is shown.

### Engineered chromosome pulling by Dis1 oligomers placed on a chromosome arm

Our experimental observations led us to postulate that a major function of kinetochores in chromosome pulling is to present Dis1 towards microtubule tips, thereby enabling stabilisation of kinetochore-microtubule attachment and efficient induction of catastrophe. This hypothesis could be tested by engineering the chromosome arm region from which a cluster of Dis1 may be presented and monitoring whether the Dis1 cluster could attach and shorten the microtubule to retrieve the chromosome arm.

The repetitive sequence of bacterial *lacO* was inserted at the *ade3* locus (approximately 2.4 Mb apart from the centromere of chromosome I) to monitor the site using the associating GFP-fused LacI protein (the *ade3::*GFP strain) ^41,42^. Endogenously expressed Dis1 was fused with the GFP-binding protein (GBP) with or without mCherry ^43,44^. Concordantly, the *ade3::*GFP foci accompanied Dis1-GBP, and were located at non-kinetochore regions, as *ade3::*GFP did not colocalise with the kinetochore marker Mis6-mTurquoise2 (*ade3::*GFP Dis1-GBP-mCherry; Supplementary Fig. 3a). The fluorescence intensity of the Dis1-GBP-mCherry oligomer was comparable to that of Dis1-mCherry (Supplementary Fig. 3b, c). This observation confirmed that Dis1 was artificially installed in the arm region of chromosomes.

The capture of artificial Dis1 oligomers assembled at the *ade3::*GFP locus by the microtubule tip significantly increased the frequency of catastrophe (I, Fig. 4a, b; Supplementary Fig. 3d; Supplementary Table 4) compared to that of free microtubules without *ade3::*GFP foci (II). Although the catastrophe frequency induced by the artificial Dis1 oligomers was moderate compared to that caused by real kinetochores (WT, Fig. 4b; Supplementary Table 4), it was sufficient to retrieve the arm region of chromosome towards SPBs in 71% of cells (Fig. 4c).

**Fig. 4:**
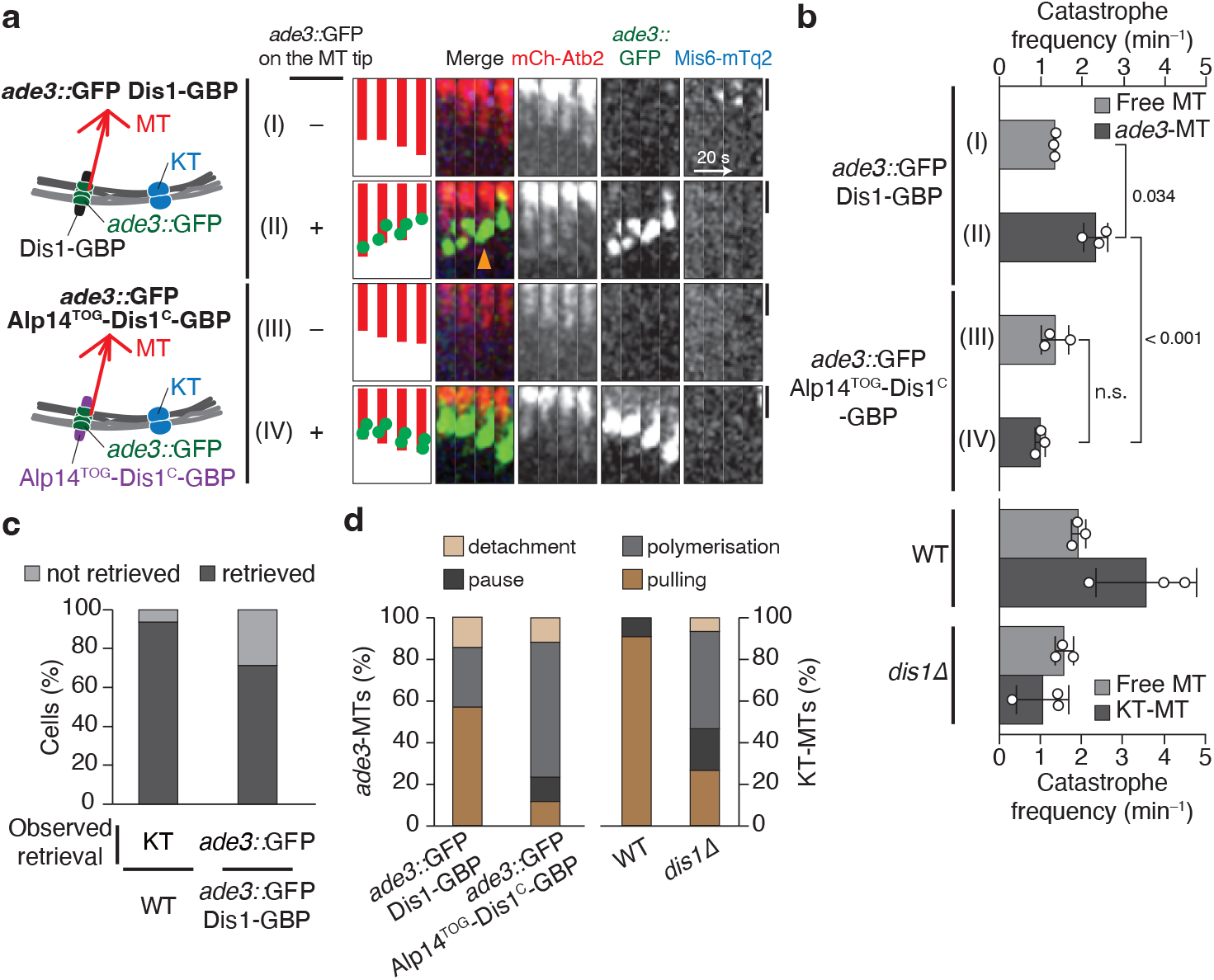
End-on pulling of chromosomes by Dis1 oligomers without using kinetochores. **a** Schematic representation of artificial pulling of chromosomes without relying on kinetochores (left). Oligomers of Dis1-GBP (top) or the chimera Alp14^TOG^-Dis1^C^-GBP (bottom) were artificially clustered at the *ade3* locus marked with GFP (*ade3::*GFP, green) on a chromosome arm. Time-lapse images of microtubules without (I and III) or with (II and IV) the *ade3::*GFP locus were filmed, and representative kymographs are shown (right). As a reference, the position of kinetochores (Mis6-mTq2, blue) is shown. The orange arrowhead indicates the start of the microtubule catastrophe. Scale bar, 1 µm. **b** The catastrophe frequency was measured for each category in (**a**). Free MT, tips without kinetochores; *ade3*-MT, tips with *ade3::*GFP. The data for WT and Dis1 are reprised from previous data as references: the data for ‘free MT’ in WT are derived from Fig. 2c (ii and iii). Other data for WT and *dis1*Δ are reprises of Fig. 2c (i, iv and v). Bullets indicate technical replicates (*n* = 3 experiments); error bars, SD. The statistical significance of the difference was determined using one-way ANOVA followed by the Tukey–Kramer method. *P* values are shown; n.s., not significant. **c** Percentages of cells that accomplished retrieval of kinetochores to SPBs in WT meiocytes (WT, *n* = 16) or retrieval of the *ade3::*GFP locus in *ade3::*GFP Dis1-GBP meiocytes (*n* = 28). χ^2^ = 3.1 (two-sample test), *P* > 0.05. **d** Frequencies of 4 events (polymerisation, pulling, pause and detachment) observed in microtubule tips accompanied by the *ade3::*GFP locus or kinetochores. *n* = 10 (WT), 13 (*dis1*Δ), 13 (*ade3::*GFP Dis1-GBP), 12 (*ade3::*GFP Alp14^TOG^-Dis1^C^-GBP) microtubules.

In WT meiocytes, the kinetochores retrieved up to the SPBs were majorly retained at this position. In contrast, 63% of engineered meiocytes detached the *ade3::*GFP foci even after reaching the SPBs (Supplementary Fig. 3e, f). This indicates that the *ade3::*GFP locus presenting Dis1-GBP was unable to maintain the arm region despite being sufficient for retrieval.

To examine whether Dis1 is solely sufficient for chromosome retrieval by microtubules, we monitored the localisation of other kinetochore factors to the artificial *ade3::*GFP site. Representative kinetochore factors such as Mis6 (centromere protein I; CENP-I) of the inner kinetochore complex (constitutive centromere associated network [CCAN]) (Fig. 4a; Supplementary Fig. 3b) as well as Ndc80 (Hec1) of the outer kinetochore network (KNL-1/Mis12 complex/Ndc80 complex [KMN]) (Supplementary Fig. 4a) were absent from *ade3::*GFP foci. The retrieval of *ade3::*GFP by Dis1-GBP oligomers was also observed in the *nuf2-2* mutant lacking the functional KMN network ^39^, demonstrating that the association of microtubule tips and the *ade3::*GFP locus via Dis1-GBP oligomers were not dependent on other kinetochore factors (Supplementary Fig. 4b–d; Supplementary Table 5).

Retrieval of the *ade3::*GFP site was not associated with the possible recruitment of the microtubule catastrophe factor kinesin-8 (Klp5-Klp6 heterodimer ^8,9,11–13^) because Klp6-3mCherry did not accumulate at the *ade3::*GFP site (Supplementary Fig. 4e). In addition, kinetochores were properly retrieved in *klp6Δ*, indicating that kinetochore retrieval does not rely on kinesin-8 in meiocytes (Supplementary Fig. 4f–h; Supplementary Table 6).

We focused on the molecular mechanism underlying Dis1-induced catastrophe. In general, TOG domains engage in the regulation of microtubules through binding to tubulin dimers ^17^. Fission yeast has two paralogous TOG proteins, unlike other eukaryotes. We, therefore, replaced the TOG domains of Dis1 with those of Alp14 and tested its function. When the chimeric protein Alp14^TOG^-Dis1^C^-GBP, comprising N-terminal TOG domains from Alp14 and C-terminal Dis1, was expressed in *ade3::*GFP cells, the chimeric oligomers localised to the *ade3::*GFP foci in a manner similar to that of the original Dis1-GBP oligomers (left, Supplementary Fig. 3a). The Alp14^TOG^-Dis1^C^-GBP chimera at *ade3::*GFP efficiently contacted microtubule tips; however, they often failed to induce microtubule catastrophe (Fig. 4a, b; Supplementary Fig. 3d; Supplementary Table 4). Dis1-GBP oligomers efficiently induced microtubules to pull the *ade3::*GFP site, whereas the chimeric oligomers failed to perform this function. In contrast, the chimeric oligomers polymerised the microtubule tips (Fig. 4d), probably reflecting the polymerase activity of Alp14 TOGs. The contrast between these two engineered oligomers mimics opposing behaviours of microtubules bound to native kinetochores observed in WT and *dis1*Δ cells (Fig. 3d).

We, therefore, concluded that the TOGs of both Alp14 and Dis1 can promote the attachment of microtubules. Nonetheless, Dis1 is exclusively responsible for the induction of microtubule catastrophe through the action of its TOG domains.

### Dis1 induces poleward motion of kinetochores in anaphase A

Our observations revealed an unconventional function of Dis1, unlike other XMAP215 orthologs, which led us to generalise that Dis1 may also direct chromosome pulling in late mitosis –– anaphase A. We monitored the segregation of sister chromatids in mitosis using the *cen2::*GFP system, in which sister centromeres of chromosome II are visualised with GFP ^41^. Sister chromatids segregated smoothly upon anaphase onset until completion of chromosome pulling, as *cen2::*GFP signals reached SPBs within 1 min (Fig. 5a). In sharp contrast, the completion of pulling took approximately 2–5 min in *dis1*Δ cells (Fig. 5a). This delay is attributed to an increase in the inter-SPB distance, as well as a frequent pause of microtubules, wherein the dynamics appeared ceased (Fig. 5a, b). These events could be interpreted as microtubule stabilisation caused by the reduction of catastrophe. Concordantly, the frequency of catastrophe that induces the transition from pause to depolymerisation was decreased in *dis1*Δ cells (Fig. 5c).

**Fig. 5:**
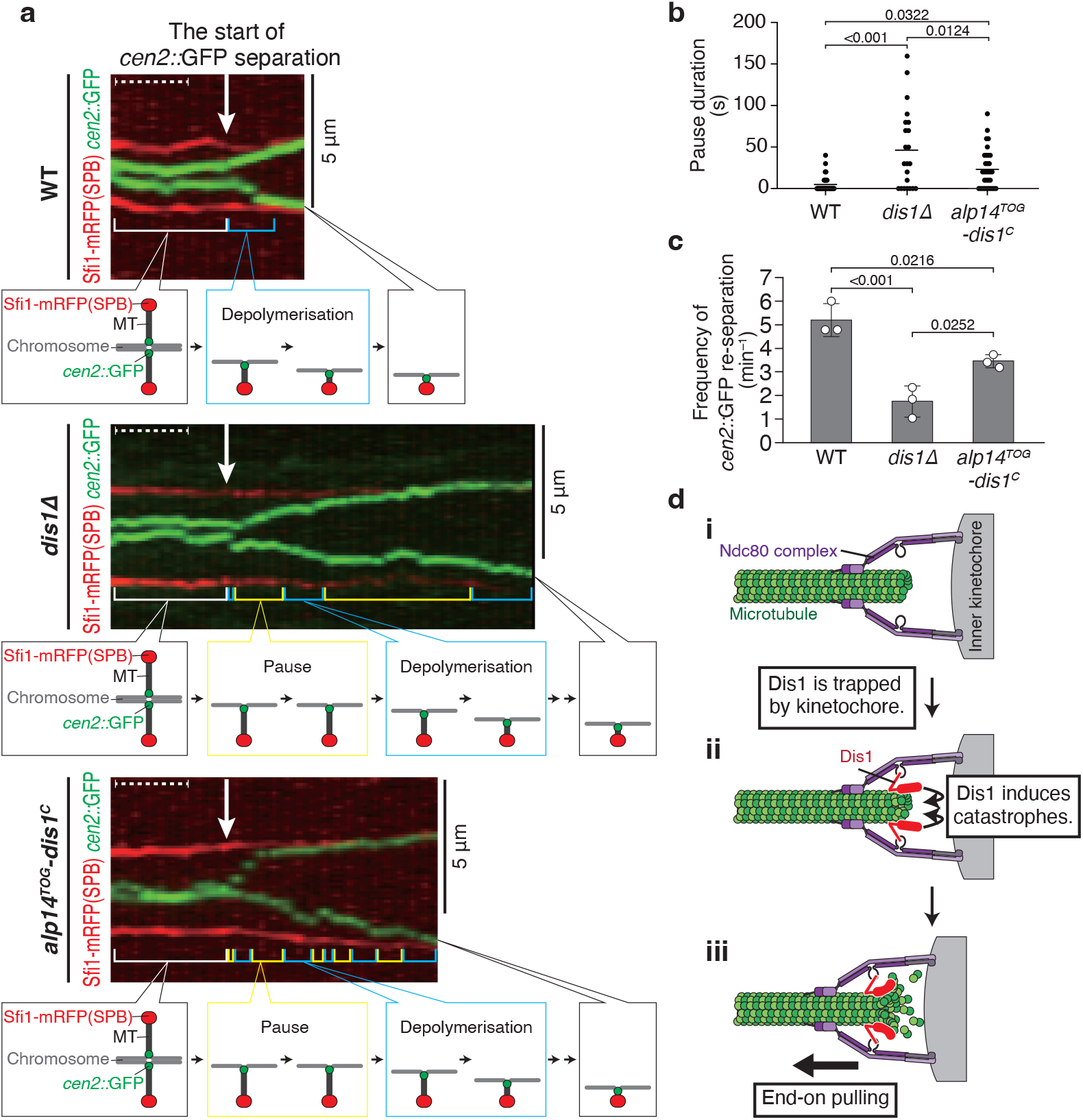
Dis1 drives kinetochore motion during Anaphase A in mitosis. **a** Representative kymographs depicting the distribution of sister centromeres of chromosome II labelled with GFP (*cen2::*GFP, green) in WT, *dis1*Δ and *alp14*^*TOG*^*-dis1*^*C*^ cells during anaphase A of mitosis. Spindle poles (SPBs) were visualised with Sfi1-mRFP (red). Arrows represent the timing when *cen2::*GFP signals started to separate. Blue brackets denote the shortening of the distance between an SPB and *cen2::*GFP, and yellow brackets denote the pause of the *cen2::*GFP signal. Dashed lines correspond to 1 min. **b** Pause duration of *cen2::*GFP in Anaphase A of each strain was plotted. *n* = 41 (WT), 14 (*dis1*Δ), 41 (*alp14*^*TOG*^*-dis1*^*C*^) centromeres. Lines represent means. **c** The frequency of *cen2::*GFP re-separation after a pause was calculated. Bullets indicate technical replicates (*n* = 3 experiments); error bars, SD. The statistical significance of the difference was determined using one-way ANOVA followed by the Tukey–Kramer method. *P* values are shown; n.s., not significant. **d** Possible schemes representing kinetochore retrieval by Dis1. First, the kinetochore attaches to the microtubule, which does not actively promote catastrophe (i). Upon reaching the microtubule tip, Dis1 is trapped by kinetochores and induces catastrophe (ii), which triggers microtubule shrinkage, thereby retrieving the kinetochore (iii).

Frequent pausing was similarly observed in the *alp14*^*TOG*^*-dis1*^*C*^ mutant, in which the TOG domain of Dis1 was replaced by that of Alp14 chimeric protein (Fig 5a–c). Although the chimeric mutant exhibited moderate defects in catastrophe induction (Fig. 5c), the double mutant of *alp14*^*TOG*^*-dis1*^*C*^ *klp6Δ* was lethal (see Discussion, Supplementary Fig. 5), suggesting that TOGs in Alp14 do not possess any adequate activities for catastrophe induction. These results collectively demonstrate that TOG domains of Dis1 are required for induction of catastrophe and are incompatible with those of Alp14.

## Discussion

Promoting catastrophe is essential in fission yeast meiocytes for the conversion of kinetochore positioning at the entry into meiosis I ^24^. Kinetochore pulling is primarily operated by the non-motor/non-kinesin protein Dis1 as a knockout of kinesin-8 (Klp5/6 heterodimers), which has been regarded as microtubule destabilizer in this organism ^8,9,11–13^, caused no apparent defects in kinetochore pulling in meiocytes. At least in meiosis, kinetochore pulling force may be mainly generated by Dis1 and supportively by kinesin-8. This study revealed the opposing activity of Dis1 to induce microtubule catastrophe *in vitro*, although TOG family members have been generally characterised or regarded as microtubule polymerase ^14,15,17–23,35^. Dis1 also induces catastrophe in mitotic cells (anaphase A) in addition to kinesin-8 (Klp5–Klp6), as supported by the fact that double knockout of *klp5* (or *klp6*) and *dis1* is synthetically lethal ^13^. The TOG domains of Dis1 are responsible for catastrophe induction during chromosome pulling in meiosis and mitosis, which are incompatible with TOGs of Alp14.

The mechanisms underlying the functional difference of Alp14 and Dis1, two TOG/XMAP215 paralogs in *S. pombe*, remain unknown. Recently, X-ray crystallography-based studies revealed that two TOG domains of Alp14 were spatially aligned in tandem, which may be crucial for Alp14 as a polymerase to attach a tubulin dimer that is eventually deposited to a microtubule tip ^45^. The AlphaFold algorithm ^46,47^ (https://alphafold.ebi.ac.uk/) was recently used to predict the similar tandem alignment of two TOGs in Alp14, whereas those in Dis1 were predicted in an antiparallel alignment with a certain degree of flexibility (schematics are shown in Supplementary Fig. 6). The flexibility of Dis1 TOGs in the structure might enable the dissociation, rather than association, of tubulin dimers from the microtubule tips, thereby possibly inducing catastrophe.

XMAP215/TOG proteins reportedly possess the activity of the microtubule polymerase; however, XMAP215 orthologs occasionally destabilise microtubules. *Xenopus* XMAP215 depolymerises GMPCPP microtubules in the egg extract ^28^ and increases both growth and shrinkage rates of microtubules ^29^. The stabilisation activity in the egg extract appears to be induced through phosphorylation ^30^. *In vitro* studies also reported the activity of *Xenopus* XMAP215 and *tobacco* MAP200 in the induction of catastrophe: this could be due to indirect effects by a simultaneous increase of the microtubule growth rate, which consequently caused an uneven extension of some protofilaments and triggered catastrophe ^48,49^. In contrast, fission yeast Dis1 appears to directly induce catastrophe during meiosis and mitosis, as microtubule growth was not enhanced during Dis1 localisation at the microtubule tips (see Figs. 2a and 4d).

In the mutants of budding yeast Stu2, the microtubule dynamics are reduced in interphase and mitosis during chromosomal attachment ^31,32^. Stu2 increased the catastrophe frequency of porcine microtubules ^33^ but decreased that of yeast microtubules ^50^ *in vitro*. Although still controversial, Stu2 may possess activities for both polymerase and catastrophe factors.

Our live-cell imaging for meiocytes demonstrated that Dis1 is mainly delivered to kinetochores via tips of growing microtubules (see Fig. 2a, Supplementary Fig. 3a), although Dis1 may stochastically be detached from the tips (i; Fig. 5d). The other TOG ortholog Alp14 is responsible for the growth of the microtubules ^24^. Once a microtubule tip reaches a kinetochore, Alp14 stabilises their attachment ^51,52^. Concomitantly, Dis1 is locked to the microtubule-kinetochore interface by the outer kinetochore factor Ndc80 (Fig. 3) ^27^, which enables continual catastrophe to shorten microtubules carrying kinetochores at the tips (ii, iii; Fig. 5d).

Artificial Dis1 oligomers are sufficient to induce microtubule catastrophe and to pull chromosome arms. Therefore, we conclude that the whole kinetochore organisation is not required for microtubule-mediated chromosomal pulling. In contrast, the role of the kinetochore is to tightly lock Dis1 to the kinetochore-microtubule interface using the Ndc80 complex ^27^. Dis1 is critical for both attachment and generation of chromosome pulling force. Notably, our experiments involving artificial retrieval of chromosome arms demonstrated that retrieved chromosomes could not be maintained by artificial Dis1 oligomers. The retention at spindle poles may require the lateral connection of microtubules and chromosomes. Dis1 might not generate sufficient force on the lateral surface of microtubules ^20^, and the outer kinetochore components represented by Ndc80 would be required for efficient retention ^36,53^. Identifying such factors would contribute to generation of an artificial system for chromosome segregation in the future.

## Methods

### Data reporting

No statistical methods were used to predetermine sample size. The experiments were not randomized. The investigators were not blinded to allocation during experiments and outcome assessment.

### Yeast strains, media and genetics

Standard materials and methods were used for *S. pombe* biology ^54^. Yeast strains used in this study are listed in Supplementary Table 7. For vegetative growth of *S. pombe* cells, YE5S (yeast extract with supplements) was used. To induce mating and meiosis, homothallic (*h*^90^) cells grown in YE5S were spotted onto sporulation agar (SPA) plates.

Standard materials and methods were used for knock-in and knock-out of genes, unless otherwise specified ^55–57^. For visualization of microtubules, coding sequences for mCherry and Atb2 (α-tubulin) were fused in frame and flanked with the native promoter and terminator for the *atb2*^*+*^ gene. The gene construct was then integrated into the Z2 region on chromosome II as an extra copy of the endogenous *atb2*^*+*^ gene, and the resultant strain is referred to as ‘*Z2-mCherry-atb2*’ in the list (Supplementary Table 7) ^58^. For tagging of the mTurquiose2 fluorescent protein to Mis6, a pFA6a-based plasmid containing the mTurquiose2 sequence and the natMX6 selection marker was created (named as pMS-mTurquoise2-nat) and used for PCR-based gene tagging as usual.

For visualisation of the position of the *ade3*^*+*^ locus on chromosome I, the canonical system utilising the bacterial lactose operator–repressor (*lacO–*LacI) was introduced ^41,42,59–61^. The *h*^90^ strain that contains the repetitive *lacO* sequences at the *ade3*^*+*^ locus (*ade3::lacO*) and expresses the fusion protein LacI-NLS-GFP (the *ade3::*GFP strain hereafter) was originally gifted by A. Yamamoto, Y. Hiraoka and M. Yamamoto.

The *ade3::*GFP strain was then genetically crossed with strains harbouring the Dis1-GBP or Dis1-GBP-mCherry fusion gene constructs ^44^. The resultant strains are expected to display Dis1-GBP or Dis1-GBP-mCherry oligomers at the *ade3* locus, respectively, in concert with LacI-NLS-GFP clustered at the *ade3::lacO* locus.

The *ndc80-21* strains with or without the *nuf2*^*+*^*-dis1(18-882)* fusion gene were gifted by T. Toda ^27^. The *nuf2-2* strain was originally gifted by Y. Hiraoka ^39^.

The strain in which the TOG domains of Dis1 were replaced with those of Alp14 (Alp14^TOG^-Dis1^C^) was constructed as follows: first, the chimeric gene construct containing the N-terminal region of Alp14 (1-500 residues) fused with the C-terminus of Dis1 (518-882), flanked by 5′- and 3′-UTR regions of the *dis1*^+^ gene, was prepared via PCR. The product was then introduced into *dis1::ura4*^+^ cells for counter-selection of *ura4*^−^ strains using YE5S plates containing 1 mg·ml^−1^ 5-fluoroorotic acid (FOA). The resultant strain *dis1::alp14*^TOG^-*dis1*^C^ therefore expresses the fusion protein Alp14^TOG^-Dis1^C^ at the endogenous level instead of Dis1. The *bsd* marker gene conferring blasticidin S resistance was inserted at the downstream of the regions of *dis1*^*+*^ gene to construct the *alp14*^*TOG*^*-dis1*^*C*^*-bsd* strain.

Centromeres on chromosome II were visualised using the previous system utilising the *cen2::*GFP system which consists of the *cen2::lacO* insertion with LacI-NLS-GFP in the *h*^90^ strain. The original *cen2::*GFP strain used herein was gifted by A. Yamamoto and Y. Hiraoka ^41^.

### Protein expression and purification

Standard methods for recombinant protein expression in bacterial cells were used, as previously summarised ^62^. To express recombinant GST-Dis1 and GST proteins in *E. coli* BL21 (DE3) cells, the coding sequence for *dis1*^*+*^ was cloned into pGEX-KG using *Sac*I and *Xba*I sites. *E. coli* cells containing plasmids were cultured in 2xYT medium at 36°C overnight. After dilution, cells were further grown for 2 hours until OD600 reaches 0.2–0.4. IPTG was then added (final 0.2 mM) for 3 h at 30°C or for 12–24 h at 20°C to induce expression. Cells were harvested by centrifugation at 5,000 rpm for 5 min, 4°C, and washed twice with PBS and PBS* (PBS with detergents and proteinase inhibitors). Cells were suspended with PBS* and lysed by sonication of repetitive ‘1-s pulse and 1-s interval’ for 1 min using the sonicator VP-50 (TAITEC). Cell debris was removed by centrifugation (5,000 rpm for 1 min at 4°C, followed by additional rounds of 14,000 rpm 1 or 3 min 4°C, ≥ 2 times) and the supernatant was collected. Glutathione Sepharose 4B (GE Healthcare Life Sciences) was added and mixed for an hour at 4°C. The Sepharose beads were then poured into a column, and washed with PBS* for 3 times. Sepharose-bound GST and GST-Dis1 proteins were eluted with the elution buffer (10 mM L-Glutathione, 50 mM Tris-HCl, pH 8.0). Eluted samples were successively fractionated into tubes. Purified protein samples were once frozen in liquid N2 and then stored at –80°C. Sample preparation was confirmed through SDS–PAGE followed by Biosafe-Coomassie (BIO-RAD) staining.

Non-labelled tubulin was purified from porcine brains by four cycles of temperature regulated polymerisation and depolymerisation in a high molarity PIPES buffer to remove contaminating MAPs ^63^. The purified tubulin was flash frozen and stored in liquid nitrogen.

### Turbidity assay

Turbidity assay was performed as previously described ^18^. Recombinant GST-Dis1 and GST were mixed on ice with 26 µM tubulin in BRB80 buffer (80 mM PIPES, 1 mM MgCl2, 1 mM EGTA, pH 6.8) containing 1 mM GTP. Microtubule polymerisation was induced by a temperature shift to 37°C, and the absorbance was monitored at 350 nm in 5-s intervals for 40 min using the spectrometer UV1800 (SHIMADZU) equipped with TCC-240A (SHIMADZU).

### Microscopy for *in vitro* assays

Microscopy regarding *in vitro* assays for tubulin dynamics was performed as previously described ^64^ with the following minor modification. Briefly, we used the inverted microscope ECLIPSE Ti (Nikon) equipped with the scanner unit CSU-W1 (Yokogawa) and the sCMOS camera Zyla 4.2 operated by the software IQ3 (Andor). To control the stage temperature, a stage-top incubator system composed of a customized double ThermoPlate chamber (11.5×7.5×0.3 cm inside size) and a TP-LH lens heater (TOKAI HIT) were used.

For visualisation of microtubules, Alexa488 (Alexa Fluor 488 NHS Ester, Thermo Fisher Scientific) labelled tubulin and unlabelled tubulin were mixed at a volume ratio of 1:19. To measure the number and length of microtubule in the presence of recombinant proteins (GST-Dis1 or GST), 22 µM pre-mixed tubulin and the recombinant proteins were mixed in BRB80 containing 0.57 mM GTP on ice. Solution was applied on a glass slide and enclosed with a cover slip (Matsunami glass) immersed with ethanol and then dry them before use. Microtubule polymerisation was induced at 37°C for 5 min, and then started observation. Fiji was used for quantification ^65^. Microtubules extending out of the image were not included.

Dynamics of microtubules was analysed as follows. First, 38 µM pre-mixed tubulin in BRB80 containing 1 mM GTP were incubated for 10 min at 37°C. The solution was then mixed with an equal volume of pre-warmed mixture comprising the observation buffer [0.8% Catalase (Sigma), 200 (U·mL^-1^) Glucose oxidase (Sigma), 9 mg·ml^−1^ Glucose, 2 mM MgCl2, 2 mM GTP, 1% (v·v^−1^) 2-mercaptoethanol] and recombinant proteins (GST-Dis1 or GST). Coverslips were immersed with ethanol and then dry them before use. Serial images were acquired every 5 s. Acquired images were converted from 16-bit to 8-bit and kymographs were generated using Fiji. In Figs. 1 and 2, images were shown in black/white inversion.

### Microscopy for cells

Our standard methods were applied as previously described ^57^. Briefly, the observation system comprised the DeltaVision-SoftWoRx image acquisition system (Applied Precision) equipped with Olympus inverted microscopes IX71 and IX81 and CoolSNAP HQ2 CCD cameras (Photometrics).

For the live-cell imaging of meiocytes, homothallic (*h*^90^) cells were spotted onto SPA and incubated for 12–13 h at 26.5°C for induction of meiosis, and additional 2 h at 32 or 36°C if necessary for temperature-sensitive strains. Cells were then mounted on a glass-bottom dish (Iwaki glass or Matsunami glass) precoated with lectin from Glycine max (Sigma). Prior to observation, the dish was filled with EMM– N+C+U+L, minimal media without a nitrogen source supplemented with uracil (50 µg·ml^−1^) and leucine (100 µg·ml^−1^) prewarmed according to the observation temperature. Images of 5–10 sections along the z-axis were acquired at 0.4-µm intervals.

For filming of mitotic cells, cells were cultivated at 30°C for overnight prior to observation in the SD medium supplemented with alanine (75 µg·ml^−1^), uracil (50 µg·ml^−1^), lysine (50 µg·ml^−1^), leucine (100 µg·ml^−1^) and histidine (50 µg·ml^−1^). Cells were mounted on a glass-bottom dish as mentioned above, and the dish was filled with EMM+N+C+5S, minimal media with a nitrogen source supplemented with alanine, uracil, lysine, leucine and histidine.

Images taken along the z-axis were deconvoluted and projected into a single image using the SoftWoRx software (v3.7.0 and v.6.5.1) and resolution was adjusted using Adobe Photoshop CC (version 2022).

The fluorescence intensity of GFP- or mCherry-tagged Dis1 in cells were quantified using SoftWoRx as follows: acquired images were projected into a single image using the Sum projection algorithm without deconvolution. A punctate signal of Dis1-GFP or Dis1-GBP-mCherry in 3 × 3 pixels was quantified using SoftWoRx, and the background signal taken outside of the nucleus was subtracted.

### Image analyses

To measure the parameters of microtubule dynamics *in vitro*, image stacks were analysed using Fiji. Then the position of the microtubule end was tracked using the MTrack J plugin for Fiji ^66^. The distance between both ends was calculated, and the transition events from microtubule growth to shrinkage were defined as catastrophe, and events from shrinkage to growth as rescue. Four parameters describing the microtubule dynamics were calculated. Growth and shrinkage rates were calculated as average values. Frequencies of catastrophe and rescue were calculated as follows: the total numbers of catastrophe events were divided by total time for microtubule growth. Kymographs were created by use of Fiji.

To measure microtubule dynamics in cells at the onset of meiosis, kymographs were created as follows: the microtubule region of interest was cropped into strips and aligned in the course of time using Adobe Illustrator CC (version 2022). The catastrophe frequency was then calculated as described above. Each section along the Z axis was analysed to confirm that the microtubule end (GFP-Atb2), Dis1-3GFP and kinetochores (marked by Mis6-mTurquiose2) are co-localised on a single plane.

The SoftWoRx software was used for tracking of *cen2::*GFP punctate signals in mitotic cells and creation of kymographs. To measure dynamic behaviour of *cen2::*GFP dots in relation to Sfi1-mCherry (an SPB marker), sequential images were applied to Fiji. The distance between an Sfi1-mCherry dot (SPB) and *cen2::*GFP (centromere) was chased over time to calculate the microtubule dynamics: pause and depolymerisation. ‘Pause’ was defined as duration without detectable motion of a *cen2::*GFP dot, whereas ‘depolymerisation’ as duration with poleward movement of *cen2::*GFP that corresponds to sister chromatid separation in anaphase A. When a paused *cen2::*GFP started to move poleward again, this depolymerisation event was particularly defined as ‘re-separation’ of *cen2::*GFP.

### Statistical analysis and reproducibility

The methods used to test for differences of statistically significant are described in each figure legend. The R package multcomp ver.1.4-19. ^67^ were used to measure of a statistical significance of difference by one-way ANOVA followed by Tukey–Kramer method. Microsoft Excel software were used to perform Student’s two tailed t-test and χ2 two-sample test. *P* values are shown on top of the corresponding columns. If the *P*-value was > 0.05, it was stated as not significant. When representative images are shown, at least three repeats were performed except for Supplementary Fig. 1a, 2a, 3a, 4a, 4e, 5. Two repeats were performed for Supplementary Fig. 2a, 3a, 4a, 4e, 5. The repeats are not necessary for Extended data Fig.1a because it represents purified proteins used in Fig. 1.

## Data availability

The data presented in this study are available from the corresponding author upon reasonable request.

## Acknowledgements

We thank A. Yamamoto, Y. Hiraoka, T. Toda and M. Yamamoto for the yeast strains. We are grateful to Y. Kakui for their valuable discussions. Y.M. was supported by JST SPRING, Grant Number JPMJSP2128. This study was supported by JSPS KAKENHI JP25291041, JP15H01359, JP16H04787, JP16H01317, JP18K19347 and JP21H00261 to M.S, and JP17K07397 and JP20K06645 to M.T. This study was also supported by the Ohsumi Frontier Science Foundation and Waseda University grants for Special Research Projects 2017B-242, 2017B-243, 2018B-222, 2019C-570, 2020R-038 and 2022C-164 to M.S and 2018S-139, 2019C-571, 2021C-584 and 2022C-170 to M.T.

## Author contributions

Y.M. conducted yeast experiments supervised by M.S. and M.T., and *in vitro* experiments supervised by T.H., M.Y. and J.Y. T.H. designed the methodologies for the *in vitro* assays. N.O. contributed to material construction. M.S. conceived the outline of the study. Y.M. and M.S. wrote the manuscript with input from the co-authors.

