## Supplementary Information for "Fission yeast Dis1 is an unconventional TOG/XMAP215 that induces microtubule catastrophe to drive chromosome pulling"

**Supplementary figures**

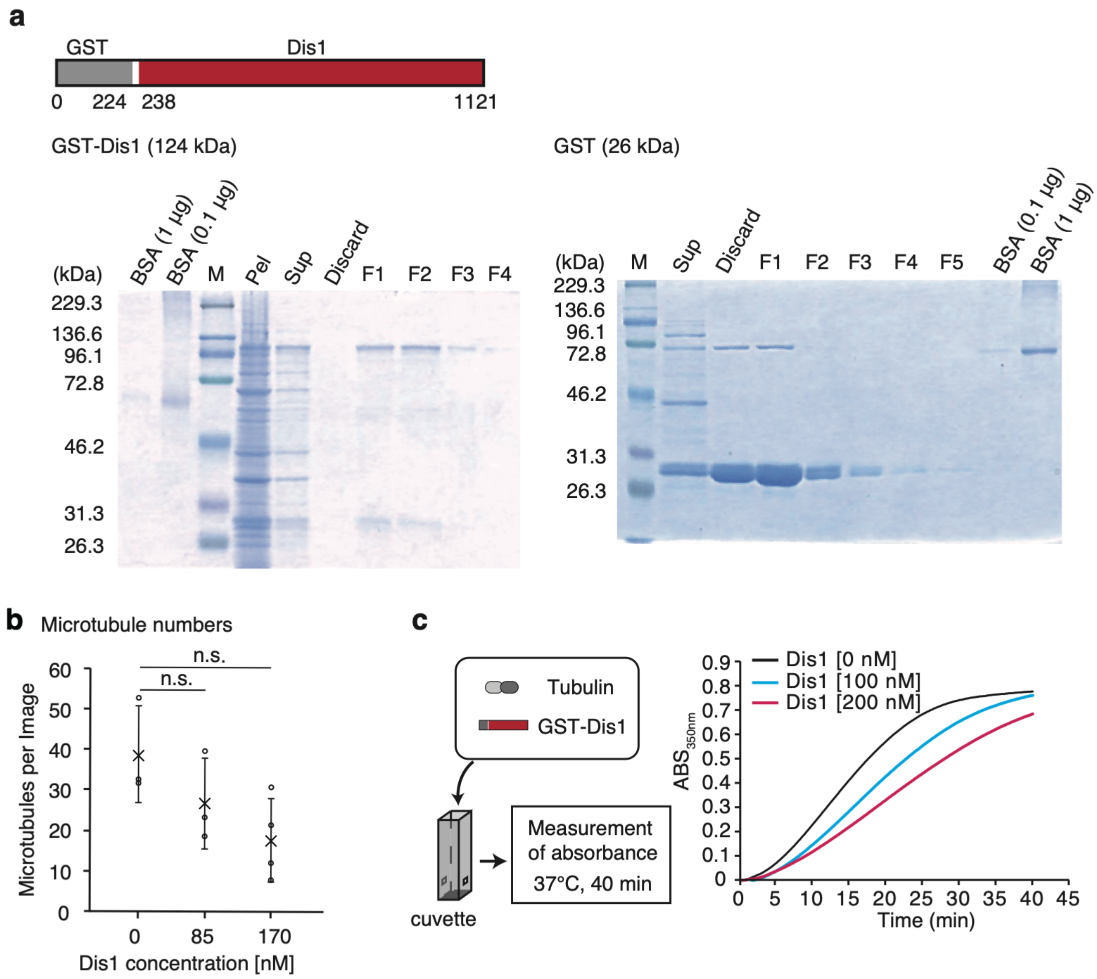


**Supplementary Fig 1. GST-Dis1 and microtubule nucleation *in vitro*.**

**a** A schematic for recombinant GST-Dis1 proteins used in this study (top). Purification of the recombinant GST-Dis1 from *E. coli* extract (bottom). Fractions of sequentially eluted samples for recombinant GST-Dis1 and GST (Discard, F1, F2, F3…) were analysed with SDS–PAGE followed by staining with Coomassie Brilliant Blue. Samples of the supernatant (Sup) and cell debris (Pel) after cell disruption were also applied. M, standards for molecular weights, shown on the left. **b** Average numbers of microtubules per observed field were plotted. The experimental procedures are as shown in **Fig. 1a**. Crosses, the mean; bullets, technical replicates ($n$ = 3 for each concentration of Dis1). More than 4 fields were analysed in each replicate. Error bars are SD. The statistical significance of difference was determined using one-way ANOVA followed by Tukey-Kramer method. $P$ values are shown; n.s., not significant. **c** Experimental outline for turbidity assays (left). Purified tubulin (26 μM) and 0–200 nM of GST-Dis1 were mixed and incubated at 37°C, and the absorbance (350 nm) was measured in 5-s intervals for 40 min. The kinetics of tubulin turbidity over time is shown as graphs. The representative of 4 technical repeats is shown.

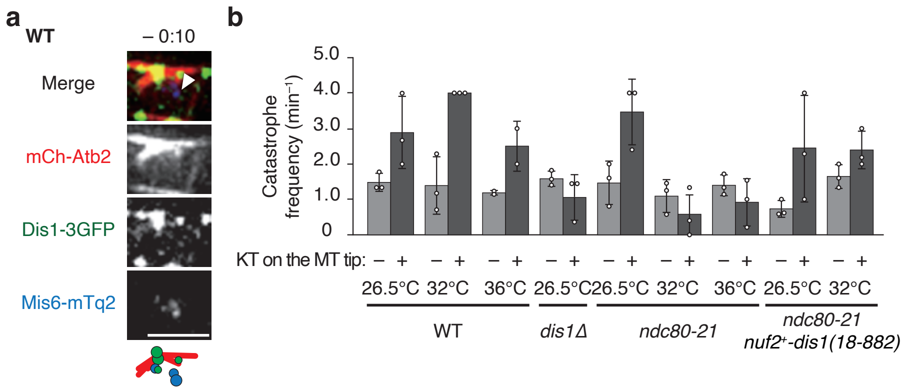


**Supplementary Fig 2. Kinetochore retrievals via microtubule catastrophe in various strains.** **a** Images for a zygotic nucleus at the onset of meiosis I in a WT cell recorded 10 s before images of 0min:00s shown in **Fig. 2a**. Dis1 was not localised to KTs that were not bound to microtubules (arrowhead). Scale bar, 2 µm. **b** Catastrophe frequencies of kinetochore-microtubules in each strain under the indicated conditions. Meiosis was induced at 26.5˚C and 32°C, similarly to **Fig. 2e**. Bullets represent 3 technical replicates, except for WT 36°C (2 replicates). Error bars, SD.

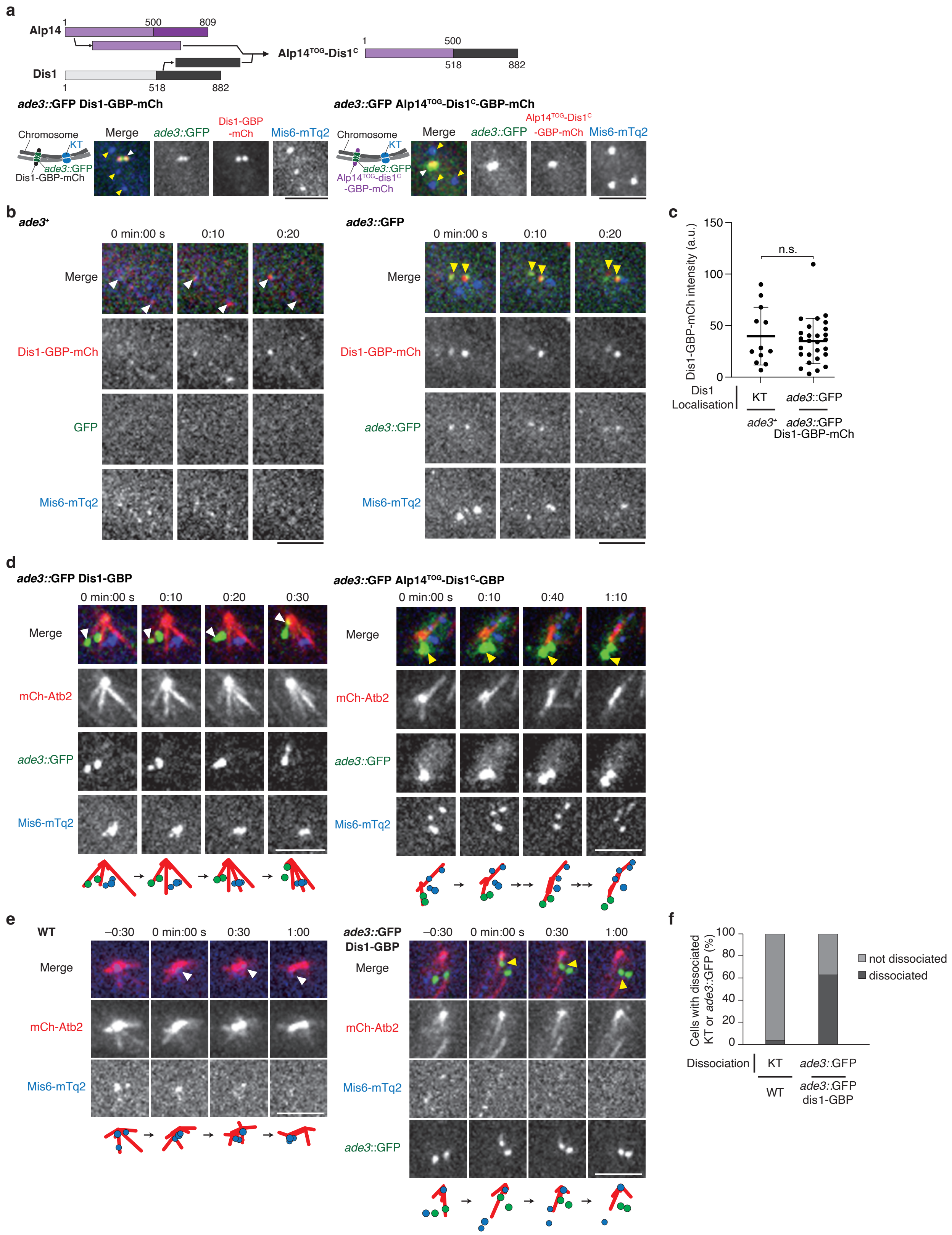


**Supplementary Fig. 3 Retrieval of the *ade3::*GFP locus with Dis1-GBP or Alp14$^{TOG}$-Dis1$^{C}$-GBP by microtubules.**

**a** A schematic for the chimeric protein Alp14$^{TOG}$-Dis1$^{C}$ used in this study (top). The *ade3* locus located on a chromosome arm region was labelled with GFP by use of LacI-GFP/LacO system. Dis1-GBP-mCherry and Alp14$^{TOG}$-Dis1$^{C}$-GBP-mCherry, and the kinetochore marker Mis6-mTurquoise2 were visualised. The GBP fusion proteins (Dis1-GBP and Alp14$^{TOG}$-Dis1$^{C}$-GBP) were colocalised with *ade3::*GFP (white arrowheads), validating that enforced oligomerisation of Dis1 (and Alp14$^{TOG}$-Dis1$^{C}$) at the *ade3::*GFP loci was successful. Note that neither Dis1-GBP-mCherry nor Alp14$^{TOG}$-Dis1$^{C}$-mCherry predominantly localised to kinetochores (marked by Mis6-mTurquoise2, yellow arrowheads). **b,c** Comparison of the Dis1 amount at kinetochores and at the *ade3::*GFP locus. In *ade3*$^{+}$ (WT) cells without GFP expression, Dis1-GBP-mCherry accumulated at kinetochores (white arrowheads, **b**). In *ade3::*GFP cells, Dis1-GBP-mCherry predominantly accumulated at *ade3::*GFP (yellow arrowheads). Signal intensities of Dis1-GBP-mCherry at kinetochores (KT, **c**) and at *ade3::*GFP were plotted. Bold lines, means. $n$ = 12 (KT) and 27 (*ade3::*GFP) foci of Dis1-GBP-mCherry. Error bars, SD. n.s., not significant (Student's two tailed t-test). **d** Time-lapse images for nuclei of *ade3::*GFP Dis1-GBP and *ade3::*GFP Alp14$^{TOG}$-Dis1$^{C}$-GBP cells at the onset of meiosis I filmed together with mCherry-Atb2 (microtubules) and Mis6-mTurquiose2 (kinetochores). The *ade3* locus was retrieved by microtubules in cells expressing Dis1-GBP (white arrowheads), but was frequently unretrieved in cells expressing the chimera-GBP (yellow arrowheads). These images were processed for kymographs shown in **Fig. 4a**. **e** Retention of retrieved chromosomes at spindle poles was monitored in WT and *ade3::*GFP Dis1-GBP nuclei. Kinetochores (Mis6-mTurquoise2) were retrieved by microtubules and retained around SPBs (white arrowheads) in WT cells, whereas the *ade3::*GFP locus was frequently dissociated from SPBs (yellow arrowheads) even once retrieved by microtubules. Scale bars, 3 μm. **f** The percentage of WT cells ($n$ = 29 cells) in which kinetochores were once retrieved to SPBs or the spindle, but dissociated from there. The percentage of *ade3::*GFP Dis1-GBP cells ($n$ = 44 cells) in which the *ade3::*GFP loci was once retrieved but dissociated. The tendency of *ade3::*GFP dissociation was statistically confirmed by the $\chi 2$ two-sample test ($\chi 2 = 22$, $P < 0.005$).

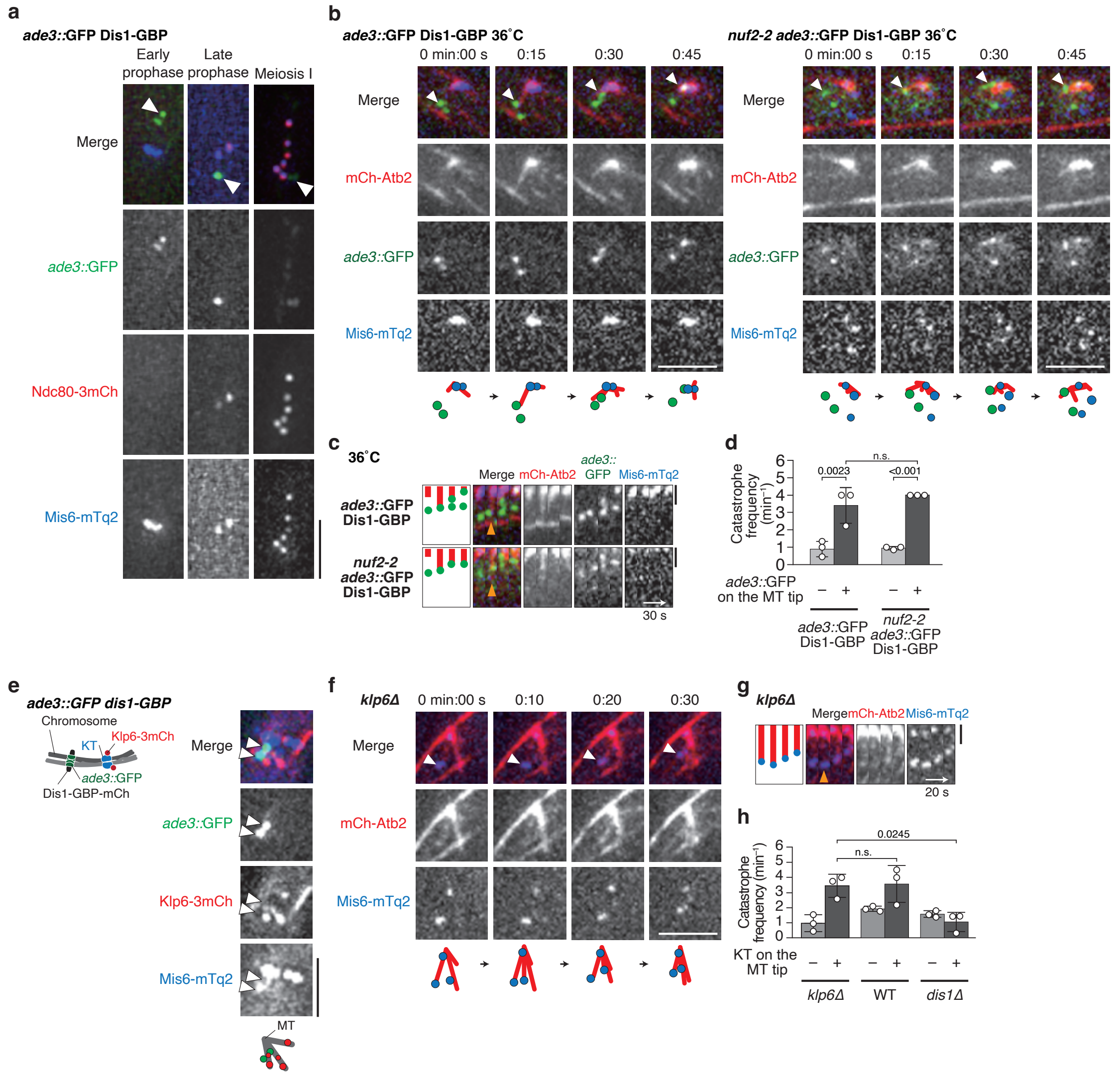


**Supplementary Fig. 4 Neither the Ndc80 complex nor Klp6 participates in microtubule shrinkage.**

**a** Images of nuclei of *ade3::*GFP Dis1-GBP cells at the indicated stages of meiosis. Ndc80-3mCherry localised nowhere (Early prophase) but to kinetochores in late prophase onwards (Mis6-mTurquoise2) [1,2]. Note that Ndc80-3mCherry was not recruited to *ade3::*GFP loci at any stage (arrowheads). **b,c** Time-lapse images of a nucleus at the onset of meiosis I in *ade3::*GFP Dis1-GBP and *nuf2-2 ade3::*GFP Dis1-GBP cells at 36°C filmed together with mCherry-Atb2 (microtubules) and Mis6-mTurquiose2 (kinetochores). In both cells, the *ade3* locus was retrieved by microtubules (arrowheads). Kymographs of the microtubules with *ade3::*GFP are also shown (**c**). The orange arrowhead indicates start of microtubule catastrophe. **d** Catastrophe frequencies of microtubules with and without *ade3::*GFP at their tips were measured for *ade3::*GFP Dis1-GBP and *nuf2-2 ade3::*GFP Dis1-GBP cells. Bullets, technical replicates: $n$ = 3 for each of four cases. **e** A meiotic nucleus with *ade3::*GFP loci expressing Dis1-GBP was filmed together with Klp6-

3mCherry and kinetochores (Mis6-mTurquoise2). Klp6-3mCherry did not co-localise with *ade3::*GFP at the onset of meiosis I (arrowheads). **f,g** Time-lapse images of a nucleus of the *klp6*Δ mutant at the onset of meiosis I. Microtubules (mCherry-Atb2) and kinetochore (Mis6-mTurquoise2) were visualised. A kinetochore retrieved by a microtubule is marked with a white arrowhead, **f.** A kymograph for the event is shown (**g**). The orange arrowhead pinpoints the start of catastrophe. Scale bars, 3 μm (**a,b,e,f**); 1 μm (**c,g**). **h** Catastrophe frequencies of microtubules with or without kinetochores in *klp6*Δ cells. Bullets, technical replicates: $n$ = 3 (+KT *klp6*Δ), 3 (–KT *klp6*Δ). The data for WT and *dis1*Δ cells are reprise of those in **Fig. 4** shown as a reference**.** Error bars, SD. The statistical significance of difference was determined using one-way ANOVA followed by Tukey-Kramer method. *P* values are shown; n.s., not significant.

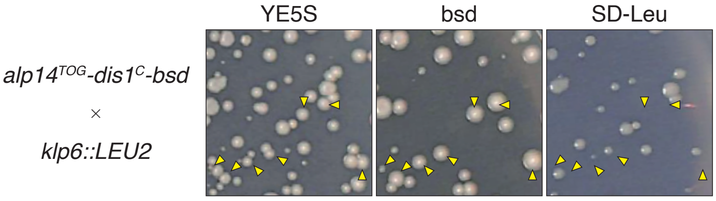


**Supplementary Fig. 5 *alp14$^{TOG}$-dis1$^{C}$* and *klp6*Δ are synthetically lethal.**

Spores from a genetic crossing of *alp14$^{TOG}$-dis1$^{C}$-bsd* and *klp6::LEU2* mutants were directly spread onto a non-selective YE5S plate for germination followed by colony formation (30°C). The colonies were then replica-plated onto YE5S plates with or without Blasticidin S (bsd) or SD–Leu plate (SD media lacking leucine) and incubated at 30°C. Colonies conferred blasticidin S resistance (arrowheads) lacked leucine autotrophy without exception, indicating that the double mutant was inviable. The *klp6* gene is located on the chromosome II, whereas *alp14$^{TOG}$-dis1$^{C}$* has been inserted to the *dis1* locus on chromosome III, therefore these two genes are not genetically linked.

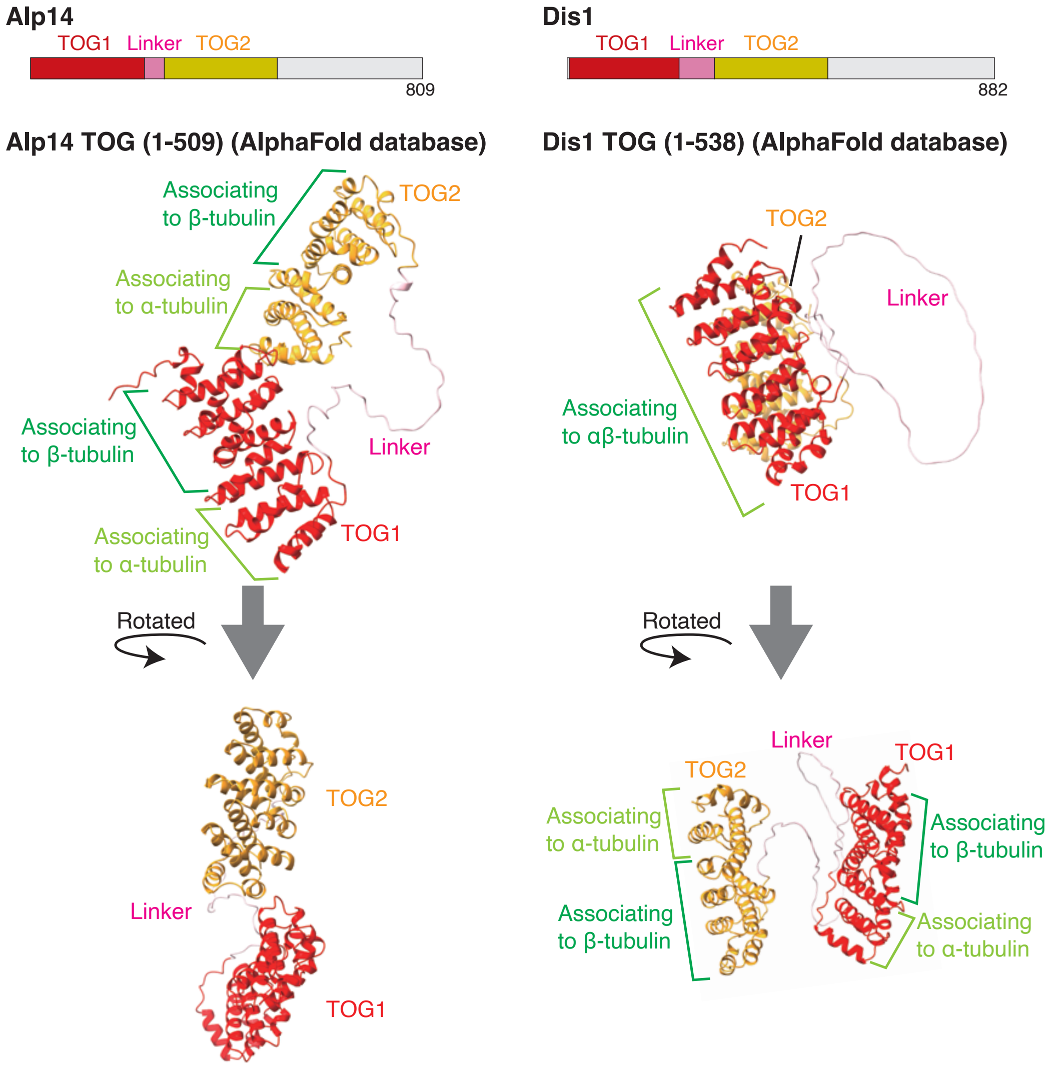


**Supplementary Fig. 6 Comparison of the predicted structures of TOG domains between Alp14 and Dis1.**

Predicted structures of the TOG domain of Alp14 (1-509 a.a.) and that of Dis1 (1-538 a.a.) were referred to the AlphaFold database (https://alphafold.ebi.ac.uk/) [3]. Domains for TOG1, linker and TOG2 are shown in red, pink and yellow, respectively, using ChimeraX v.1.2.5. (UCSF) [4]. The style of association between tubulin dimers and each TOG was adapted from a previous study on Alp14 crystallography [5].

**Supplementary tables**

**Supplementary Table 1. Parameters of microtubule dynamics *in vitro***

| Dis1 concentration [nM] | The number of microtubules (sum) | Catastrophe frequency (min–1) | Rescue frequency ($min^{-1}$) | Growth rate ($\mu m \cdot min^{-1}$) | Shrinkage rate ($\mu m \cdot min^{-1}$) |
|---|---|---|---|---|---|
| 0 | 95 | 0.073 ± 0.020 | 0.10 ± 0.20 | 0.98 ± 0.19 | 22.5 ± 4.59 |
| 50 | 125 | 0.114 ± 0.023 | 0.80 ± 0.26 | 1.19 ± 0.08 | 19.8 ± 1.98 |
| 100 | 148 | 0.116 ± 0.010 | 1.46 ± 0.66 | 1.20 ± 0.11 | 17.4 ± 2.96 |
| 200 | 162 | 0.118 ± 0.020 | 1.52 ± 0.56 | 1.44 ± 0.13 | 15.4 ± 3.88 |

Frequencies of catastrophe and rescue, and rates for growth and shrinkage are mean ± SD (*n* = 4 experiments). See methods for details.

**Supplementary Table 2. Parameters of microtubule dynamics in wild-type and *dis1*Δ cells**

| Genotype | Existence of Dis1 and KT on the microtubule tip | The total number of microtubules (sum) | Catastrophe frequency ($min^{-1}$) |
|---|---|---|---|
| WT | –Dis1–KT | 22 | 1.7 ± 0.36 |
| WT | +Dis1–KT | 17 | 2.4 ± 0.20 |
| WT | +Dis1+KT | 11 | 3.6 ± 1.22 |
| *dis1*Δ | *dis1*Δ –KT | 52 | 1.6 ± 0.22 |
| *dis1*Δ | *dis1*Δ +KT | 13 | 1.1 ± 0.64 |

Catastrophe frequencies of microtubules at meiosis I onset in strains of each genotype were calculated. Observed microtubules were classified depending on whether each microtubule displayed Dis1 and kinetochores (KT) on the tip. The total number of microtubules observed in 3 experiments are also shown. The frequencies are mean ± SD (3 experiments).

**Supplementary Table 3. Parameters of microtubule dynamics in wild-type, *ndc80-21* and *ndc80-21 nuf2$^{+}$-dis1(18-882)* cells**

| Genotype | Existence of KT on the microtubule tip | Temperature | The total number of microtubules (sum) | Catastrophe frequency ($min^{-1}$) |
|---|---|---|---|---|
| WT | –KT | 26.5°C | 22 | 1.5 ± 0.26 |
| WT | +KT | 26.5°C | 9 | 2.9 ± 1.0 |

| | | | | |
|---|---|---|---|---|
| WT | –KT | 32°C | 11 | 1.4 ± 0.81 |
| WT | +KT | 32°C | 8 | 4.0 ± 0 |
| WT | –KT | 36°C | 7 | 1.2 ± 0.062 |
| WT | +KT | 36°C | 6 | 2.5 ± 0.71 |
| *ndc80-21* | –KT | 26.5°C | 9 | 1.5 ± 0.61 |
| *ndc80-21* | +KT | 26.5°C | 7 | 3.5 ± 0.92 |
| *ndc80-21* | –KT | 32°C | 18 | 1.1 ± 0.46 |
| *ndc80-21* | +KT | 32°C | 17 | 0.58 ± 0.68 |
| *ndc80-21* | –KT | 36°C | 8 | 1.4 ± 0.29 |
| *ndc80-21* | +KT | 36°C | 10 | 0.92 ± 0.67 |
| *ndc80-21 nuf2*$^+$*-dis1(18-882)* | –KT | 26.5°C | 15 | 0.73 ± 0.23 |
| *ndc80-21 nuf2*$^+$*-dis1(18-882)* | +KT | 26.5°C | 9 | 2.4 ± 1.5 |
| *ndc80-21 nuf2*$^+$*-dis1(18-882)* | –KT | 32°C | 6 | 1.6 ± 0.34 |
| *ndc80-21 nuf2*$^+$*-dis1(18-882)* | +KT | 32°C | 11 | 2.4 ± 0.53 |

Catastrophe frequencies of microtubules at meiosis I onset were calculated for each strain with the indicate genotype at the indicated temperature. Observed microtubules were classified depending on whether each microtubule displayed kinetochores (KT) on the tip. The total number of microtubules observed in 2 experiments (WT, 36˚C) or 3 experiments (others) are also shown. The frequencies are mean ± SD (2 experiments for WT, 36˚C; 3 experiments for others).

**Supplementary Table 4. Parameters of microtubule dynamics in *ade3::*GFP Dis1-GBP and *ade3::*GFP Alp14$^{TOG}$-Dis1$^{C}$-GBP cells**

| Genotype | Existence of *ade3::*GFP on the microtubule tip | The total number of microtubules (sum) | Catastrophe frequency (min$^{-1}$) |
|---|---|---|---|
| *ade3::*GFP Dis1-GBP | –*ade3::*GFP | 30 | 1.3 ± 0.027 |
| *ade3::*GFP Dis1-GBP | +*ade3::*GFP | 14 | 2.3 ± 0.29 |

| | | | |
|---|---|---|---|
| *ade3::*GFP Alp14$^{TOG}$-Dis1$^{C}$-GBP | –*ade3::*GFP | 26 | 1.3 ± 0.33 |
| *ade3::*GFP Alp14$^{TOG}$-Dis1$^{C}$-GBP | +*ade3::*GFP | 17 | 0.98 ± 0.12 |

Catastrophe frequencies of microtubules at meiosis I onset in strains of each genotype were calculated. Observed microtubules were classified depending on whether each microtubule displayed *ade3::*GFP foci on the tip. The total number of microtubules observed in 3 experiments are also shown. The frequencies are mean ± SD (3 experiments).

**Supplementary Table 5 Parameters of microtubule dynamics in *ade3::*GFP Dis1-GBP and *nuf2-2 ade3::*GFP Alp14$^{TOG}$-Dis1$^{C}$-GBP cells**

| Genotype | Existence of *ade3::*GFP on the microtubule tip | The total number of microtubules (sum) | Catastrophe frequency (min$^{-1}$) |
|---|---|---|---|
| *ade3::*GFP Dis1-GBP | –*ade3::*GFP | 9 | 0.89 ± 0.44 |
| *ade3::*GFP Dis1-GBP | +*ade3::*GFP | 14 | 3.4 ± 1.0 |
| *nuf2-2* *ade3::*GFP Dis1-GBP | –*ade3::*GFP | 6 | 0.95 ± 0.091 |
| *nuf2-2* *ade3::*GFP Dis1-GBP | +*ade3::*GFP | 9 | 4.0 ± 0.00 |

Catastrophe frequencies of microtubules at meiosis I onset in strains of each genotype were calculated. Observed microtubules were classified depending on whether each microtubule displayed *ade3::*GFP foci on the tip. The total number of microtubules observed in 3 experiments are also shown. The frequencies are mean ± SD (3 experiments).

**Supplementary Table 6 Parameters of microtubule dynamics in *klp6*Δ cells**

| Genotype | Existence of KT on the microtubule tip | The total number of microtubules (sum) | Catastrophe frequency (min$^{-1}$) |
|---|---|---|---|
| *klp6*Δ | –KT | 31 | 0.97 ± 0.55 |
| *klp6*Δ | +KT | 14 | 3.4 ± 0.76 |

Catastrophe frequencies of microtubules at meiosis I onset in *klp6*Δ cells were calculated.

Observed microtubules were classified depending on whether each microtubule displayed Dis1 and kinetochores (KT) on the tip. The total number of microtubules observed in 3 experiments are also shown. The frequencies are mean ± SD (3 experiments).

**Supplementary Table 7 *S. pombe* strains used in this study**

| Strain | Alias | Genotype | Origin | Related figures |
|---|---|---|---|---|
| YM0439 | Wild type | *h90 dis1-3GFP-kan Z2-mCherry-atb2-hph mis6-mTurquoise2-nat leu1 ura4 ade6-M216* | This study | Figure 2a, 2b, 2c, 3a, 3b, 3c, 4c, 4d, S2a, S3e, S3f |
| YM0451 | *dis1Δ* | *h90 dis1::ura4+ Z2-mCherry-atb2-hph mis6-mTurquoise2-nat leu1 ura4 ade6-M216* | This study | Figure 2a, 2b, 2c, 4d |
| YM0455 | Wild type | *h90 Z2-mCherry-atb2-hph mis6-mTurquoise2-nat leu1 ura4 ade6-M216* | This study | 2d, 2e, S3b |
| YM0512 | *ndc80-21* | *h90 ndc80-21-kan Z2-mCherry-atb2-hph mis6-mTurquoise2-nat leu1 ura4 ade6-M216* | This study | Figure 2d, 2e, S3b |
| YM0540 | *ndc80-21 $nuf2^{+}$-dis1(18-882)* | *h90 ndc80-21-bsd nuf2+-dis1(18-882)-ura4+ Z2-mCherry-atb2-hph mis6-mTurquoise2-nat leu1 ura4 ade6-M216* | This study | Figure 2d, 2e, S3b |
| YM0567 | *ade3::GFP dis1-GBP* | *h90 dis1-GBP-bsd ade3::LacO-ura4+-kan his7+-LacI-GFP Z2-mCherry-atb2-hph mis6-mTurquoise2-nat leu1 ura4 ade6-M216* | This study | Figure 4a, 4b, 4c, 4d, S3d, S3e, S3f, S4b, S4c, S4d |
| YM0604 | *ade3::GFP $alp14^{TOG}$-$dis1^{C}$-GBP* | *h90 dis1::$alp14^{TOG}$-$dis1^{C}$-GBP-bsd ade3::LacO-ura4+-kan his7+-LacI-GFP Z2-mCherry-atb2-hph mis6-mTurquoise2-nat leu1 ura4 ade6-M216* | This study | Figure 4a, 4b, 4d, S3d |
| MJ0018 | Wild type | *h90 sfi1-mRFP-hph cen2-LacO-kan-ura4+ his7+-LacI-GFP leu1 ura4 lys1 ade6-M216* | Our stock | Figure 5 |

| | | | | |
|---|---|---|---|---|
| RD0025 | *dis1Δ* | *h90 dis1::bsd sfi1-mRFP-hph cen2-LacO-kan-ura4+ his7+-LacI-GFP leu1 ura4 ade6-M216* | Our stock | Figure 5 |
| YM0918 | *alp14*$^{TOG}$*-dis1*$^{C}$ | *h90 dis1::alp14*$^{TOG}$*-dis1*$^{C}$ *sfi1-mRFP-hph cen2-LacO-kan-ura4+ his7+-LacI-GFP leu1 ura4 ade6-M216* | This study | Figure 5 |
| YM0586 | *ade3::GFP dis1-GBP-mCh* | *h90 dis1-GBP-mCherry-hph ade3::LacO-ura4+-kan his7+-LacI-GFP mis6-mTurquoise2-nat leu1 ura4 ade6-M216* | This study | Figure S3a, S3b, S3c |
| YM0620 | *ade3::GFP alp14*$^{TOG}$*-dis1*$^{C}$*-GBP-mCh* | *h90 dis1::alp14*$^{TOG}$*-dis1*$^{C}$*-GBP-mCherry-hph ade3::LacO-ura4+-kan his7+-LacI-GFP mis6-mTurquoise2-nat leu1 ura4 ade6-M216* | This study | Figure S3a |
| YM0924 | Wild type | *h90 dis1-GBP-mCherry-hph mis6-mTurquoise2-nat leu1 ura4 ade6-M216* | This study | Figure S3b, S3c |
| YM0634 | *ade3::GFP dis1-GBP ndc80-3mCh* | *h90 ndc80-3mCherry-hph dis1-GBP-bsd ade3::LacO-ura4+-kan his7+-LacI-GFP mis6-mTurquoise2-nat leu1 ura4 ade6-M216* | This study | Figure S4a |
| YM0932 | *nuf2-2 ade3::GFP Dis1-GBP* | *h90 nuf2-2::ura4+ dis1-GBP-bsd ade3::LacO-ura4+-kan his7+-LacI-GFP mis6-mTurquoise2-nat leu1 ura4 ade6-M216* | This study | Figure S4b, S4c, S4d |
| YM0635 | *ade3::GFP dis1-GBP klp6-3mCh* | *h90 klp6-3mCherry-hph dis1-GBP-bsd ade3::LacO-ura4+-kan his7+-LacI-GFP mis6-mTurquoise2-nat leu1 ura4 ade6-M216* | This study | Figure S4e |
| YM0925 | *klp6Δ* | *h90 klp6::LEU2 Z2-mCherry-atb2-hph mis6-mTurquoise2-nat leu1 ura4* | This study | Figure S4f, S4g, S4h |
| YM0936 | *alp14*$^{TOG}$*-dis1*$^{C}$*-bsd* | *h90 alp14*$^{TOG}$*-dis1*$^{C}$*-bsd leu1 ura4* | This study | Figure S5 |
| LJ0272 | *klp6Δ* | *h90 klp6::LEU2 leu1 ura4* | Our stock | Figure S5 |
